# The UIP honeycomb airway cells are the site of mucin biogenesis with deranged cilia

**DOI:** 10.1101/2022.09.03.506451

**Authors:** Jeremy A. Herrera, Lewis A. Dingle, M. Angeles Montero, Rajamiyer V. Venkateswaran, John F. Blaikley, Felice Granato, Stella Pearson, Craig Lawless, David J. Thornton

**Author notes:** Corresponding author, University of Manchester, 4.019 AV Hill, Oxford Road, Manchester, Greater Manchester, M13 9PT, United Kingdom.

## Abstract

Honeycombing (HC) is a histological pattern consistent with Usual Interstitial Pneumonia (UIP). HC refers to cystic airways (HC airways) located at sites of dense fibrosis with marked mucus accumulation. Utilizing laser capture microdissection coupled mass spectrometry (LCM-MS), we interrogated the fibrotic HC airway cells and fibrotic uninvolved airway cells (distant from sites of UIP and morphologically intact) in 10 UIP specimens; 6 non-fibrotic airway cell specimens served as controls. Furthermore, we performed LCM-MS on the mucus plugs found in 6 UIP and 6 mucinous adenocarcinoma (MA) specimens. The mass spectrometry data were subject to both qualitative and quantitative analysis and validated by immunohistochemistry. Surprisingly, fibrotic uninvolved airway cells share a similar protein profile to HC airway cells, showing deregulation of SLITs and ROBO pathway as the strongest category. We find that BPIFB1 is the most significantly increased secretome-associated protein in UIP, whereas MUC5AC is the most significantly increased in MA. We conclude that spatial proteomics demonstrates that the fibrotic uninvolved airway cells are abnormal. In addition, fibrotic HC airway cells are enriched in mucin biogenesis proteins with a marked derangement in proteins essential for ciliogenesis. This unbiased spatial proteomic approach will generate novel and testable hypotheses to decipher fibrosis progression.

## Introduction

Usual Interstitial Pneumonia (UIP) is a fibrotic disease that is associated with a variety of fibrotic entities (Idiopathic Pulmonary Fibrosis – IPF, sarcoidosis, non-specific interstitial pneumonia - NSIP, connective tissue disease - CTD, and hypersensitivity pneumonitis - HP) [1–3]. The UIP histological pattern is patchy with regions of relatively normal-appearing lung adjacent to dense fibrosis and honeycombing (HC). HC refers to the clustering of airspaces within dense fibrotic tissue and is associated with the thickening of airway walls. Accumulation and plugging of mucus and other airway debris within the HC airways likely negatively impacts lung function.

Our current understanding of fibrotic airway pathogenesis has been improved with the advancement of structural, genetic, and molecular analyses. Structurally, UIP/IPF lung have reduced numbers of terminal bronchioles in both regions of minimal and established fibrosis [4–6]. Genetically, sequence changes affecting alveolar cells (*MUC5B, SFTPC*, and *SFTPA2*) have been reported [7–9]. Cells comprising the HC airways present as either multi-layer or as a single layer [1], with cellular subtypes including; basal, ciliated, columnar, pseudostratified and secretory epithelium; while there are variable reports on the presence of alveolar type II (ATII) cells [10–12]. Functionally, single-cell RNA sequencing of IPF epithelial cells identify marked cellular heterogeneity as compared to control [13]. Collectively, these factors are believed to lead to airway homeostasis impairment and facilitate disease progression.

An important function of the airway is to produce mucus. Not only does mucus serve as a physical barrier but mucus has antimicrobial properties to protect distant airways [14]. During the fibrotic process, mucus fills and plugs the HC airways, which affects pathogen clearance and blood-oxygen exchange. The secreted mucus hydrogel is underpinned by two gel-forming mucins, of which, mucin 5B (MUC5B) is the most abundant in health, whereas MUC5AC is also detected but at a lower level [10]. A gain-of-function *MUC5B* polymorphism is amongst one of the highest risk factors associated with lung fibrosis [10, 15–18]. However, knowledge on the molecular composition of the mucus plug in UIP is incomplete.

We have recently created a tissue atlas of the fibrotic front of UIP/IPF utilizing laser capture microdissection coupled mass spectrometry (LCM-MS) [19, 20]. In this study, we used the same approach to define the composition and provide mechanistic themes of the fibrotic HC airway cells with the aim to identify key targets and pathways to intercept fibrosis progression. In addition, we identify the composition of mucus in HC airways in lung fibrosis (UIP) and compare this to lung cancer (mucinous adenocarcinoma [MA]) to determine if mucus heterogeneity exists in varying disease states where mucus plugs are found in the airways.

## Results

### Laser capture microdissection of fibrotic and non-fibrotic airway cells

Figure 1. shows our approach to laser capture microdissection (LCM). Using alcian blue/periodic acid Schiff’s (AB/PAS) stain, mucus is visualized as purple in color within the fibrotic HC airway (**Figure 1A, upper row**); note how the AB/PAS stain lines the airway cells (red arrows) in a manner that suggests mucin is being secreted centrally into the airway lumen. We show that we precisely captured the mucus in a fibrotic specimen, including its cellular infiltrates (**Figure 1A, middle row**). In addition, we captured the mucin-rich epithelial lining of HC airways (**Figure 1A, lowest row**). We also captured fibrotic uninvolved airway cells defined as being in distant regions demonstrating minimal fibrosis (**Figure 1B**). Our LCM capabilities allow us to precisely isolate this region while leaving behind mucus associated in uninvolved airways, denoted with a red asterisk. To serve as a control, we performed LCM on airway cells from non-fibrotic specimens (**Figure 1C**). In total, we performed LCM on 10 fibrotic specimens (n = 10 fibrotic HC airway cells and n = 10 fibrotic uninvolved airway cells) and on 6 non-fibrotic airway cell controls (a total of 26 samples).

**Figure 1:**
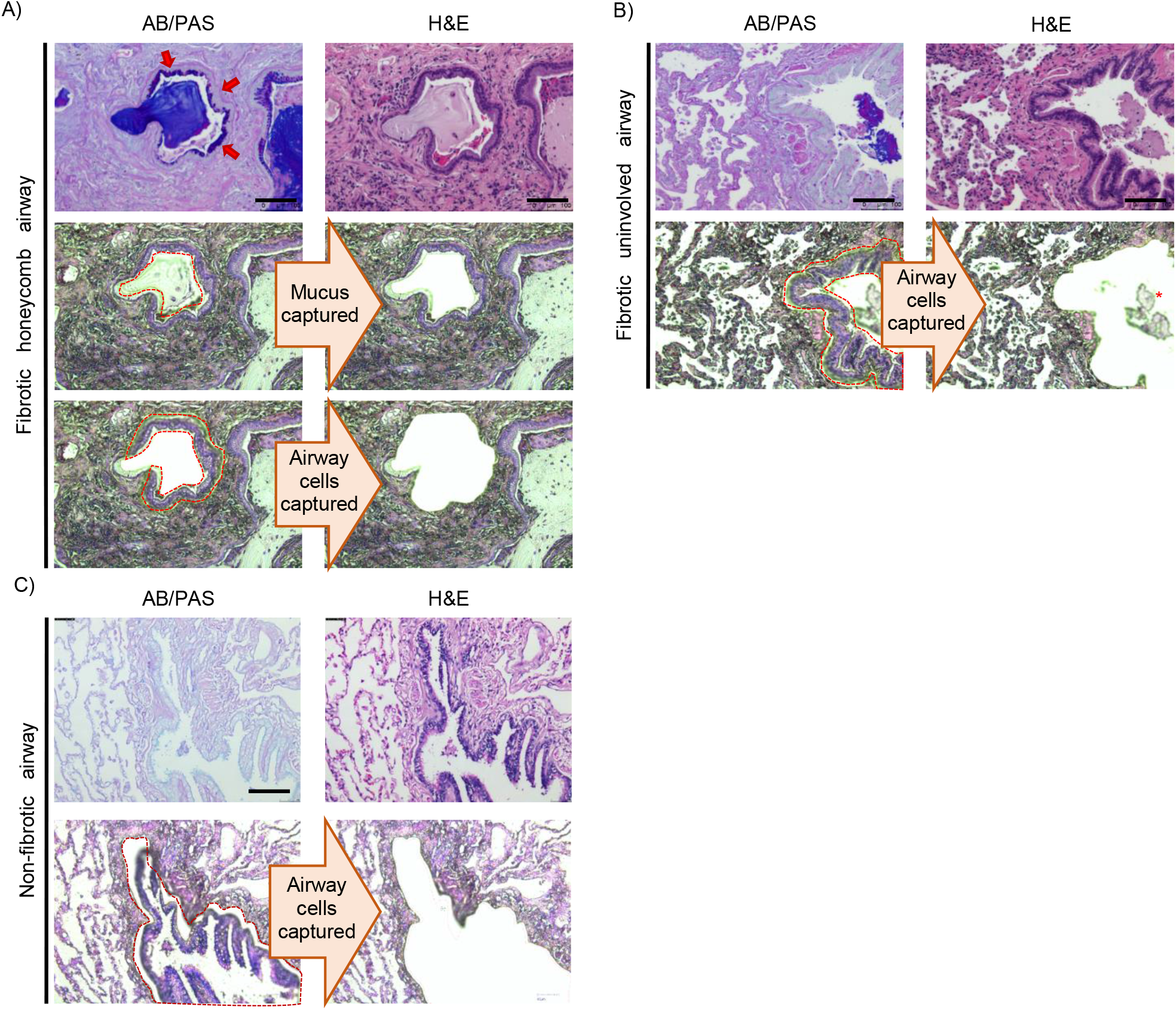
Laser capture microdissection of the mucus, fibrotic honeycomb, fibrotic uninvolved and non-fibrotic airway cell controls. Formalin-fixed paraffin-embedded specimens were serially sectioned at 5 microns and stained with alcian blue/periodic acid Schiff’s (AB/PAS) stain or Hemetoxylin & Eosin (H&E). (**A**) A representation of laser capture microdissection in a fibrotic specimen. AB/PAS (Top left) or H&E (the other 5 panels). Mucus (purple) was visualized with AB/PAS stain; notice how the mucin lines the inner airway consistent with these cells producing mucin centrally into the airspace [red arrows]. We individually captured the mucus and fibrotic honeycomb airway cells for mass spectrometry preparation. (**B**) In the same fibrotic patient, we found uninvolved airways in the morphologically intact regions of the fibrotic lung and captured the airway cells for mass spectrometry preparation (notice how the mucus plug is left behind– depicted with red asterisk). (**C**) A representation of non-fibrotic airway cells captured for mass spectrometry preparation. Scale bar represents 100 microns.

### The fibrotic HC and uninvolved airway cells are similar in protein composition

We prepared our samples for mass spectrometry (MS) following our established protocol [19, 20] and performed a qualitative analysis to determine which proteins are present per group: non-fibrotic, fibrotic uninvolved, and fibrotic HC airway cells. We define a protein present if it is detected in 3 of the 6 non-fibrotic airway cell samples or 5 of the 10 fibrotic airway cell samples. We detected 2,668 proteins in human lung airway cells (**Figure 2A**) and provide a complete list of these proteins (**Supplemental File 1**). We found that more proteins are detected in fibrotic HC airway cells, which may be attributed to the metabolically demanding process of mucin production [21].

**Figure 2.**
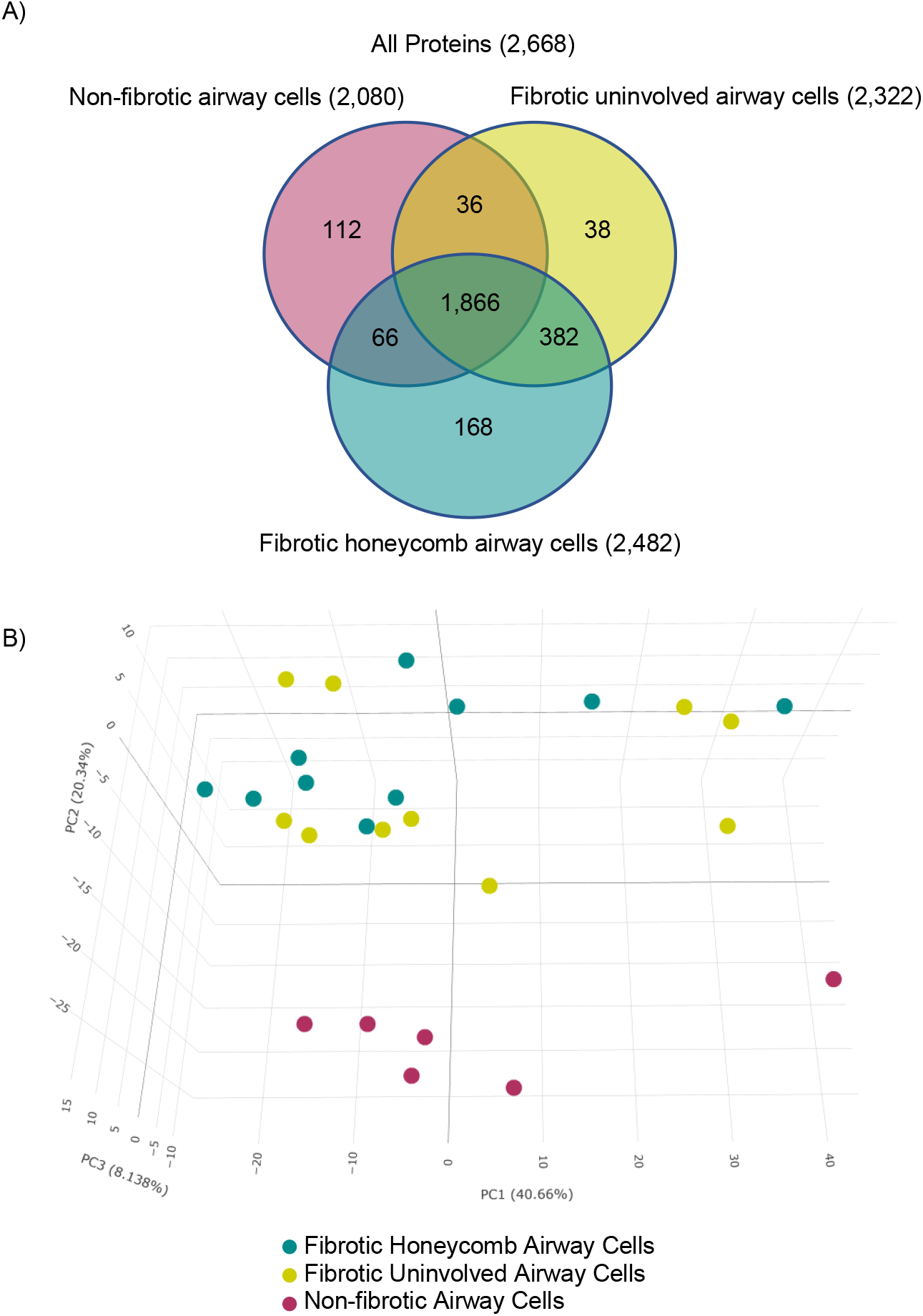
Spatial proteomic analysis of the fibrotic airway cells. Fibrotic and non-fibrotic specimens were subjected to laser capture microdissection coupled mass spectrometry (LCM-MS) to collect non-fibrotic airway cells (n = 6 specimens), fibrotic uninvolved airway cells (n = 10 specimens), and fibrotic honeycomb (HC) airway cells (n = 10 specimens). (**A**) Venn diagram showing the number of proteins found in each airway cell type. (**B**) 3-D Principal component analysis showing that the non-fibrotic airway cells (red dots) cluster away from the fibrotic airway cells (the other dots). Surprisingly, fibrotic uninvolved airway cells (yellow dots) and HC airway cells (green dots) cluster together.

We next performed a quantitative analysis, which compares the relative abundance of the detected proteins, to create a 3-dimensional (3-D) principal component analysis (PCA) (**Figure 2B**). Firstly, we showed that non-fibrotic airway cells (red dots) separate from both the fibrotic HC (green dots) and fibrotic uninvolved airway cells (yellow dots). Surprisingly, we found that both the fibrotic HC airway cells and fibrotic uninvolved airway cells closely cluster with some deviation. This analysis suggests that the fibrotic uninvolved airway cells (found in morphologically intact lung) display an abnormal protein profile as the mucin-rich HC airway cells (found within the densely fibrotic region of the lung).

### Honeycomb airway cells are enriched in proteins involved in mucus biogenesis and have decreased cilia-associated proteins

Although the fibrotic HC and fibrotic uninvolved airway cells cluster by PCA, we sought to further compare these groups. Of the 2,957 proteins detected, we found that there are 101 proteins significantly increased in fibrotic HC airway cells while 18 are statistically increased in the fibrotic uninvolved airway cells (**Figure 3A**, a full list in **Supplemental File 2**). A list of the highest and lowest proteins is provided in **Table 1**. Consistent with our approach of capturing mucin-rich HC airway cells, we found that MUC5B is significantly increased in the fibrotic HC airway cells and not in the fibrotic uninvolved airway cells.

**Table 1:** Highest and lowest 15 significantly changed proteins in the fibrotic HC airway cells compared to fibrotic uninvolved airway cells.

| ***Increased in fibrotic HC airway cells*** |  |  | ***Decreased in fibrotic HC airway cells*** |  |  |
| --- | --- | --- | --- | --- | --- |
| ***Protein*** | ***Log<sub>2</sub>*** | ***FDR*** | ***Protein*** | ***Log<sub>2</sub>*** | ***FDR*** |
| BPIFB1 | 3.97 | 4.60E-24 | CEP135 | -1.30 | 3.18E-02 |
| DHRS9 | 3.79 | 1.40E-03 | COL8A1 | -1.23 | 3.28E-03 |
| MUC5B | 3.65 | 1.70E-21 | FBLN2 | -1.00 | 2.16E-02 |
| S100P | 2.77 | 2.40E-19 | LRRC45 | -0.90 | 2.88E-02 |
| FAM3D | 2.73 | 2.90E-02 | TTC21B | -0.86 | 3.29E-03 |
| BPIFA1 | 2.52 | 7.40E-04 | LAMB2 | -0.75 | 3.40E-04 |
| LCN2 | 2.20 | 3.13E-13 | APCS | -0.74 | 4.45E-02 |
| CRABP2 | 2.08 | 3.22E-02 | CGN | -0.74 | 1.45E-02 |
| DMBT1 | 1.74 | 4.82E-03 | FBN1 | -0.72 | 2.24E-02 |
| CPD | 1.68 | 2.30E-06 | IFT57 | -0.72 | 4.30E-05 |
| ST6GAL1 | 1.64 | 1.02E-02 | NCEH1 | -0.69 | 6.30E-04 |
| IGJ | 1.55 | 3.763E-03 | PLG | -0.63 | 8.00E-03 |
| AGR2 | 1.53 | 1.00E-05 | H1F0 | -0.62 | 6.25E-03 |
| FKBP11 | 1.49 | 1.53E-02 | CYP51A1 | -0.60 | 2.90E-02 |
| SLPI | 1.46 | 1.80E-04 | CERS2 | -0.59 | 2.59E-02 |

**Figure 3:**
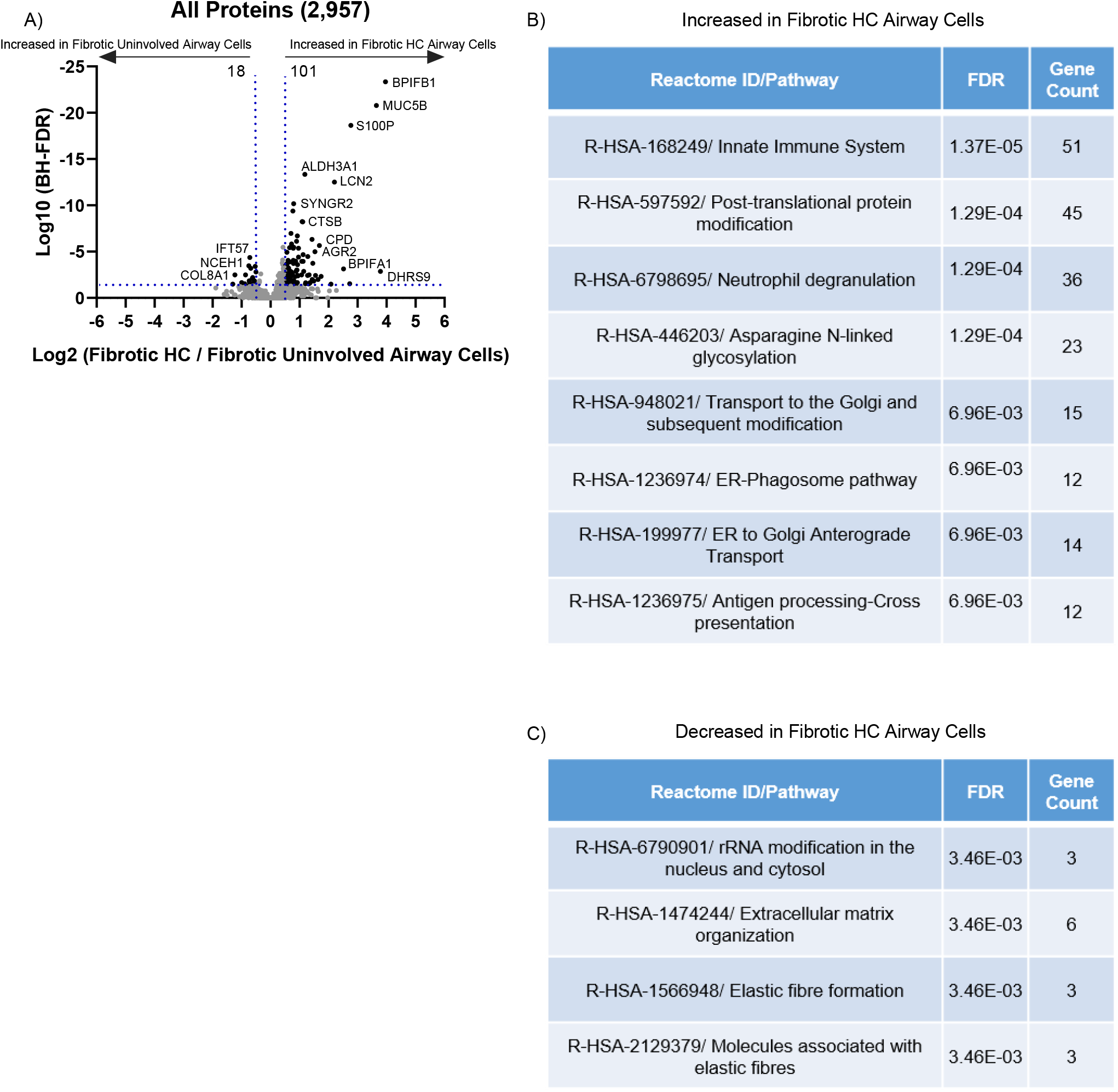
The fibrotic honeycomb airway cells have a pro-mucin protein signature. (**A**) Volcano plot comparing the fibrotic honeycomb (HC) airway to fibrotic uninvolved airway cells showing the negative natural log of the false discovery values (FDR) values plotted against the base 2 log (fold change) for each protein. Reactome pathway showing the most (**B**) increased or (**C**) decreased for the fibrotic HC airway cells compared to fibrotic uninvolved airway cells.

Strikingly, many of the proteins increased in the fibrotic HC airway cells are involved in mucin biogenesis and/or regulation. For instance, bactericidal/permeability-increasing (BPI) fold-containing family B member 1 (*BPIFB1*) is a negative regulator of *MUC5B* expression [22] and is at the top of the list. Similarly, secretory leukocyte protease inhibitor (*SLPI*) reduces mucin expression *in vitro* and is enriched in the fibrotic HC airway cells [23]. This suggests that negative regulators of mucin expression in lung fibrosis are insufficient to stop mucin production. Retinoic acid signalling induces mucin gene expression and secretion [24, 25]; Dehydrogenase reductase SDR family member 9 (*DHRS9*) and cellular retinoic acid-binding protein 2 (*CRABP2*) both modulate retinoic acid synthesis and are increased in the fibrotic HC airway cells. Recently, CRABP2 was shown to be increased in IPF airway cells [26]. In accord with mucin biogenesis, anterior gradient homolog 2 (*AGR2*) has been shown to be essential for MUC2 production and FK506-binding protein 11 (*FKBP11*) has been demonstrated to have a mucin secretory function [27]. Both AGR2 and FKBP11 are increased in the fibrotic HC airway cells. Some unique proteins to the fibrotic HC airway cells are GALNT12, GALNT3, ST6GAL1, and GALNT6, which are associated with O-linked glycosylation of mucins. In accord with these findings, Reactome pathway analysis demonstrated that a variety of pathways pertaining to mucin production are increased, such as ‘Post-translational protein modification’, ‘Transport to the Golgi and subsequent modification’, and ‘ER-phagosome pathway’ (**Figure 3B**); unfortunately, mucin biogenesis is not an established Reactome pathway. Reactome pathway analysis also demonstrated that ‘extracellular matrix organization’ and ‘elastic fibre formation’ are decreased in the fibrotic HC airway cells [**Figure 3C**]. We confirm that there are disorganized elastic fibres in the HC airways, which is in accord with the loss of airway structure and increased fibrosis in this region (**Supplemental Figure 1**). Thus, our spatial proteomic approach identifies a pro-mucin protein signature associated with the fibrotic HC airway cells as compared to fibrotic uninvolved airway cells.

**Supplemental Figure 1.**
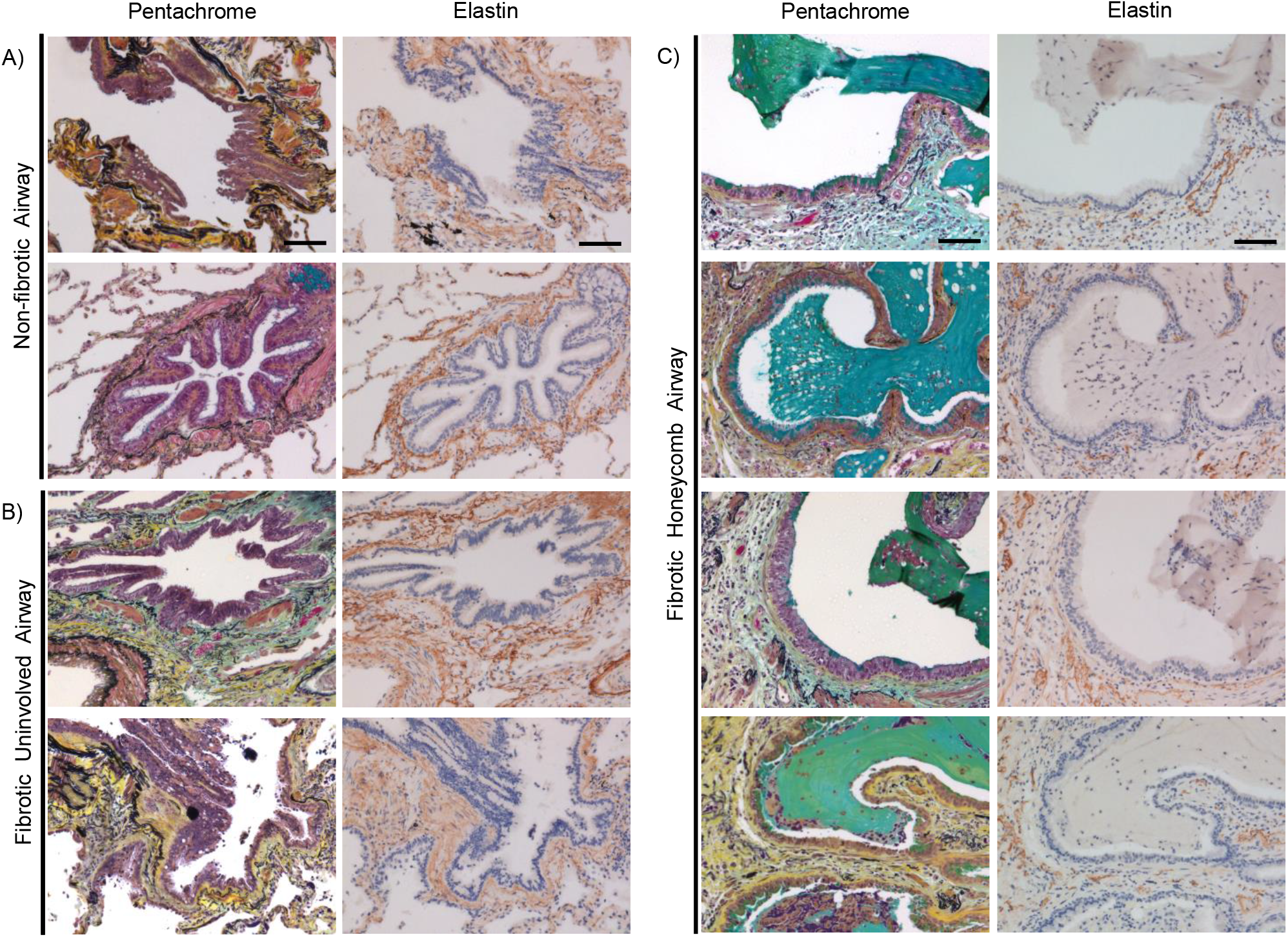
Elastin disorganization in the fibrotic honeycomb airway. 2 Non-fibrotic and 4 fibrotic specimens were stained for pentachrome or immunostained for elastin. Shown are representative images for (**A**) non-fibrotic airway, (**B**) fibrotic uninvolved airways, and (**C**) fibrotic honeycomb airways. Note that elastin fibres (black in color in the pentachrome) surround airways in the non-fibrotic airway and fibrotic uninvolved airways, but is disorganized in the honeycomb airways.

Cilia are conserved organelles that function to clear airway mucus and associated debris. Herein, we demonstrated that multiple proteins associated with ciliogenesis are decreased in the fibrotic HC airway cells. For instance, Centrosomal protein 135 (CEP135) is the most decreased protein and is required for ciliogenesis initiation [28]. Ciliogenesis relies on a variety of proteins, including intraflagellar transport (IFT) proteins [29]. Intraflagellar transport protein 57 (IFT57) is required for cilia maintenance and is decreased in the fibrotic HC airway cells; other intraflagellar transport proteins (IFT81, IFT46) are not expressed in the fibrotic HC airway cells but expressed in the non-fibrotic or fibrotic uninvolved airway cells. Leucine-rich repeat protein 21B (*LRRC45*) and tetratricopeptide repeat protein 21B (*TTC21B*) are critical for ciliogenesis and are also decreased in the fibrotic HC airway cells [30, 31]. A variety of proteins that are not expressed in the fibrotic HC airway cells include proteins associated with cilia function, such as CEP131, CCP110, KIF3A, CYB5D1, DYNLRBR2, RPGR, and WRD66 [32–38]. These data suggest that the loss of proteins associated with ciliogenesis within the fibrotic HC airways may be part of the mechanism of fibrosis progression.

A previous study demonstrated that ciliary microtubules are morphologically disorganized in UIP/IPF, which will have profound effects on cilia structure and function [39]. To demonstrate that abnormal ciliogenesis is a potential mechanistic theme in the fibrotic HC airway cells, we immunostained for tubulin alpha 4a (TUBA4*A*; a marker of cilia) in 4 UIP specimens and 2 controls. We found that the cilia marker is widely expressed in cells lining the airway cells of non-fibrotic and fibrotic uninvolved airways (**Figure 4A – 4B**). In contrast, the mucin-rich regions of the HC airway (red arrows) are largely devoid of cilia (black arrows) (**Figure 4C**).

**Figure 4.**
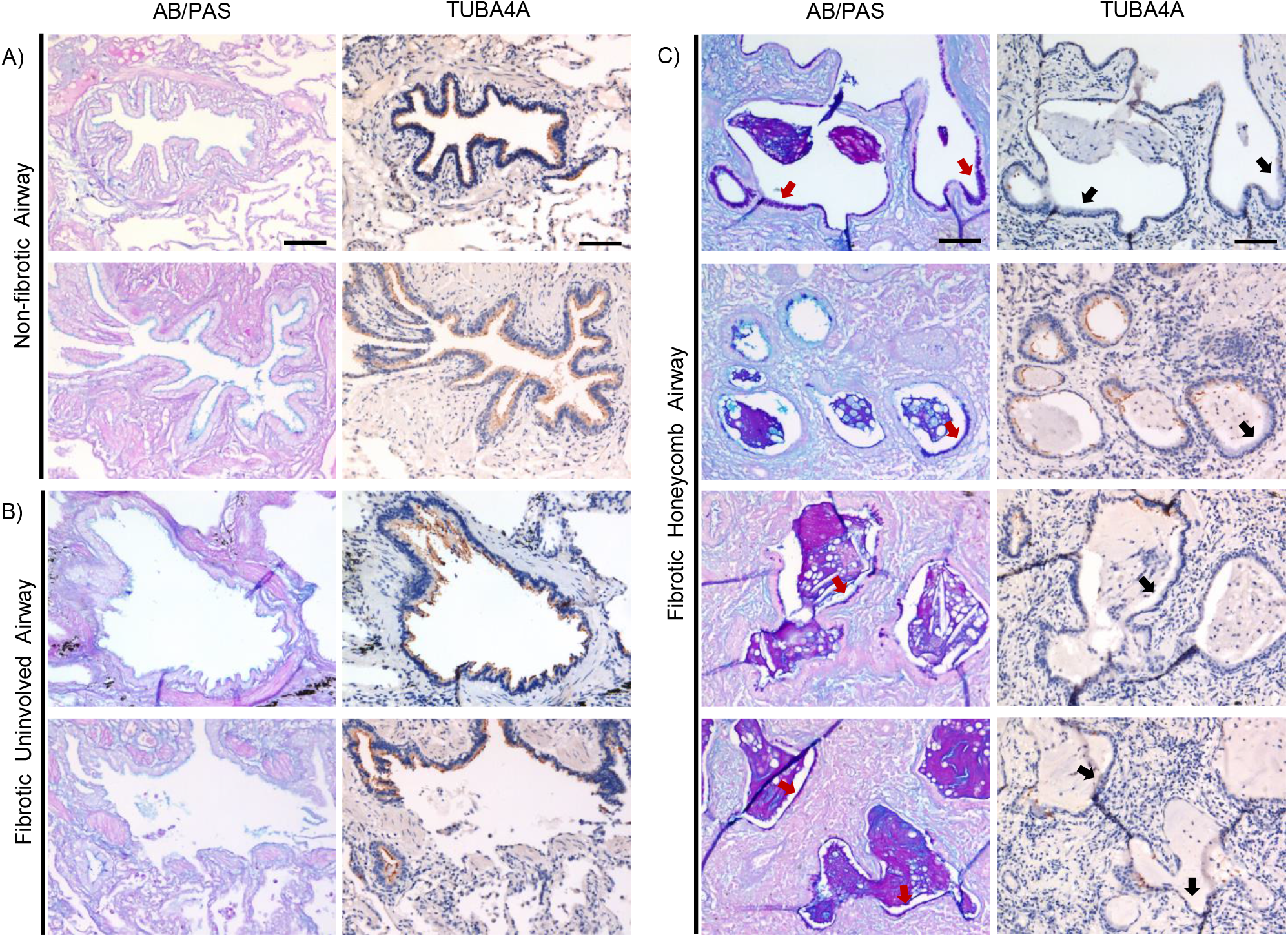
Cilia expression in the fibrotic honeycomb airway cells. Two Non-fibrotic and 4 fibrotic specimens were stained for alcian blue/periodic acid Schiff’s (AB/PAS) or immunostained for TUBA4A (a marker of cilia). Shown are representative images for (**A**) non-fibrotic airway, (**B**) fibrotic uninvolved airways, and (**C**) fibrotic honeycomb (HC) airways. Note that regions of mucin positivity (red arrows) are absent of cilia (black arrows) in the fibrotic HC airway cells. Scale bar represents 100 microns.

### The fibrotic uninvolved airway cells have an abnormal protein signature

We next compared both the fibrotic airway cells to the non-fibrotic airway cell controls. We showed that 333 proteins are significantly increased in the fibrotic HC airway cells, whereas 157 proteins are significantly increased in the non-fibrotic airway cell controls (**Figure 5A**; a full list in **Supplemental File 2**). Reactome pathway analysis demonstrated that ‘regulation of expression of *SLIT*s and *ROBO*s’ and ‘Signaling by *ROBO* receptors’ are the strongest categories increased in the fibrotic HC airway cells as compared to non-fibrotic controls (**Figure 5B**). The Slit/Robo pathway is implicated in a variety of cellular processes, including neurogenesis, cell proliferation, and migration [40]. Slit/Robo has been shown to be involved in airway development [41, 42], and can improve alveolar regeneration in lung-injury models [43]. We next compared the fibrotic uninvolved airway cells to the non-fibrotic airway cell controls. We detected 178 proteins significantly increased in fibrotic uninvolved airway cells, whereas we found 202 proteins significantly increased in non-fibrotic airway cell controls (**Figure 5C**; a full list in **Supplemental file 2**). Surprisingly, the 15 highest proteins increased in the fibrotic uninvolved airway cells are also increased in the fibrotic HC airway cells. Reactome pathway analysis again show that ‘Regulation of expression of *SLIT*s and *ROBO*s’ and ‘Signaling by *ROBO* receptors’ are the strongest categories in the fibrotic uninvolved airway cells as compared to non-fibrotic controls (**Figure 5D**); Reactome pathway analysis did not find any significantly decreased pathway enrichment in either fibrotic airway groups. Given that similar pathways and proteins are increased in the fibrotic uninvolved airway cells as the fibrotic HC airway cells, suggests that the fibrotic uninvolved airway cells are abnormal.

**figure 5:**
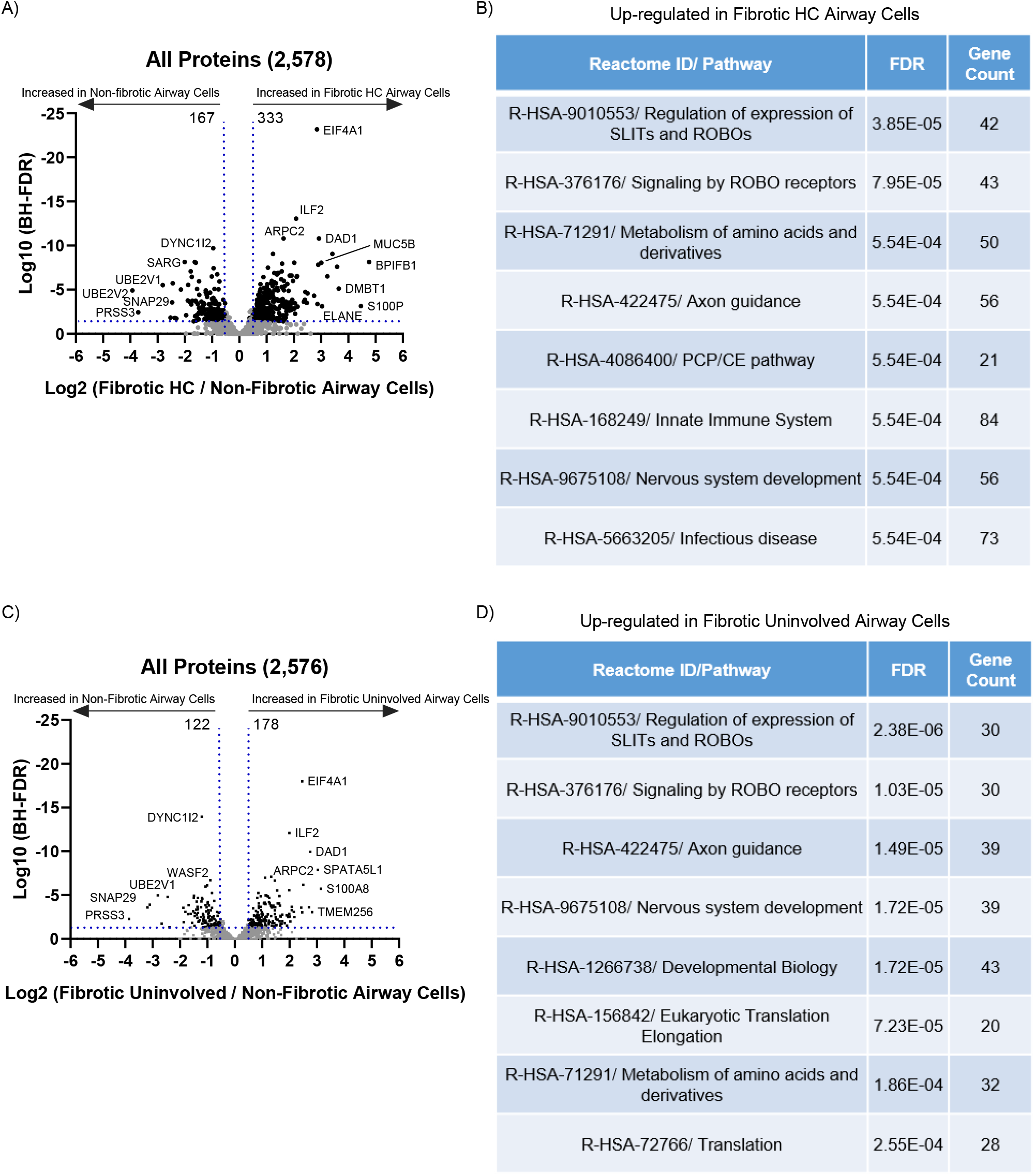
The fibrotic uninvolved airway cells have an abnormal protein signature. (**A**) Volcano plot omparing the fibrotic honeycomb airway cells to non-fibrotic airway cells showing the negative natural log of the lse discovery values (FDR) values plotted against the base 2 log (fold change) for each protein. (**B**) Reactome athway showing the most increased pathway for the fibrotic honeycomb airway cells compared to non-fibrotic irway cells. (**C**) Volcano plot comparing the fibrotic uninvolved airway cells to non-fibrotic airway cells showing e negative natural log of the FDR values plotted against the base 2 log (fold change) for each protein. (**D**) eactome pathway showing the most increased pathway for the fibrotic uninvolved airway cells compared to on-fibrotic airway cell controls.

A heatmap of the 568 significantly changed proteins across the groups: fibrotic HC, fibrotic uninvolved, and non-fibrotic airway cell controls is shown in **Figure 6A** (a full list in **Supplemental File 3**). The fibrotic HC airway cells share similarities to the fibrotic uninvolved airway cells, but with substantial deviations. We next show the 25 highest and lowest changed proteins (**Figure 6B**). Ras-related protein 3D (RAB3D) was the most increased in the fibrotic HC airway cells and is involved in the biogenesis of secretory granules [44]. MUC5B and MUC5AC are both packaged in the secretory granules of airway cells [45]. Similarly, prolyl endopeptidase (PREP) is found within exosomes in airway cells and released upon LPS stimulation [46]. Cyclase associated protein 1 (CAP1) is associated with lung cancer and post-translational modification promotes proliferation and migration [47]. At the bottom of the list are dynein axonemal assembly factor 2 (DNAAF2), matrix gla protein (MGP), and agrin (AGRN) which are only decreased in the fibrotic HC airway cells (detected in the fibrotic uninvolved and non-fibrotic airway cell controls). *DNAAF2* is involved in cilia homeostasis and mutations in *DNAAF2* lead to cilia defects [48]. *MGP* is considered an inhibitor of calcification based on the extensive cardiovascular calcification observed in *MGP*-null mice [49]; calcification occurs in UIP/IPF patients, and is associated within regions of honeycombing [50]. *AGRN* is a proteoglycan that serves a variety of biological functions, including the promotion of regeneration [51]. Further work interrogating the collective roles of these changed proteins will help decipher the mechanism of fibrosis progression.

**Figure 6:**
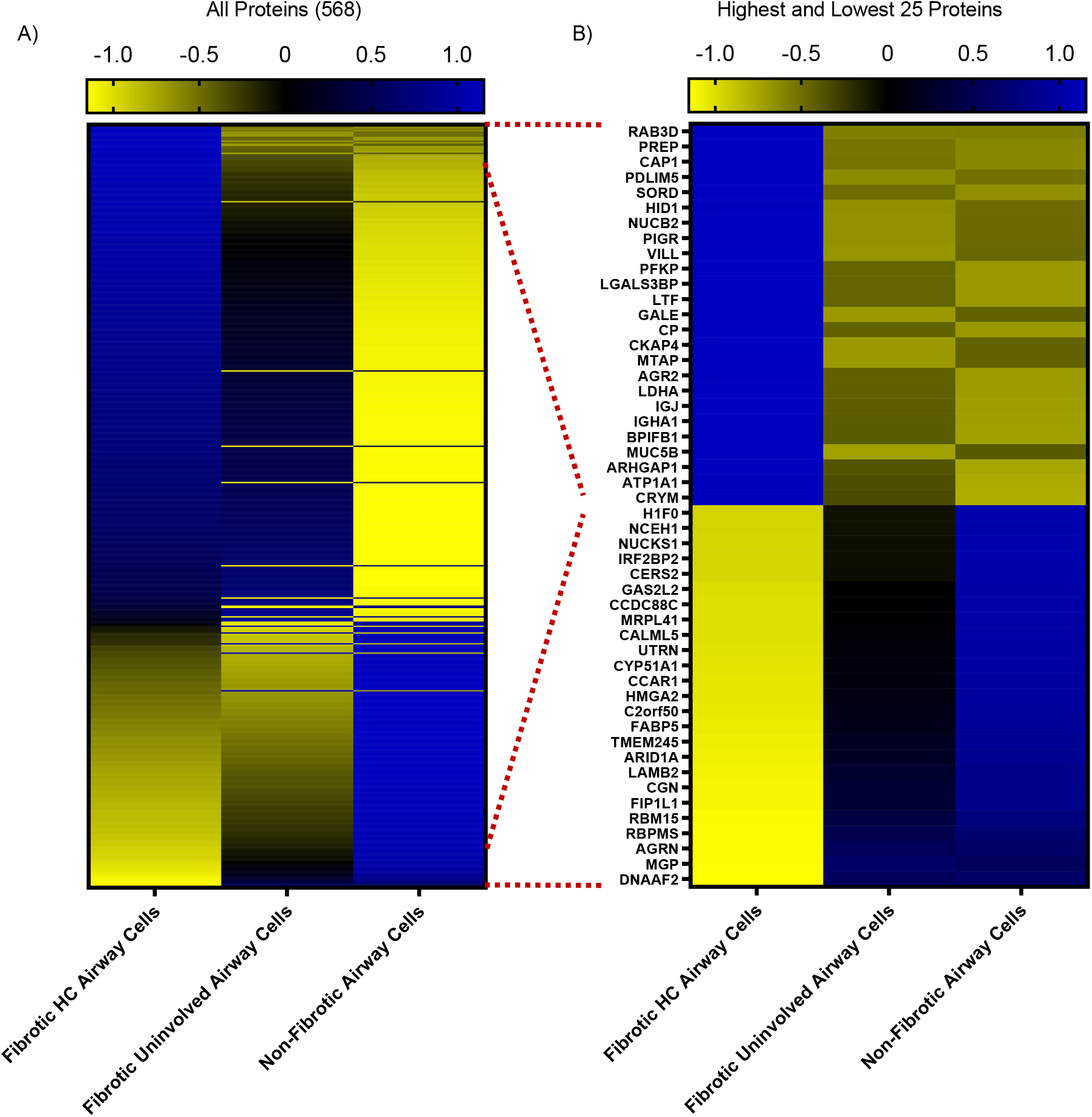
Fibrotic airway cells differ than controls. Shown are heatmaps of (**A**) all 568 statistically changed proteins or (**B**) the highest and lowest 25 proteins in the fibrotic honeycomb, fibrotic uninvolved, and non-fibrotic airway cells. Proteins are arranged by increasing abundance with reference to the honeycomb airway cells.

### The composition of fibrotic lung mucus

We previously demonstrated our capacity to microdissect the mucus plugs in fibrotic specimens (**Figure 1A**, middle panels). Utilizing our MS approach, we detected 650 proteins in the fibrotic/UIP mucus plugs (detected in 3 or more of the 6 samples; **Supplemental File 4**). Using intensity Based Absolute Quantification (iBAQ; a measure of protein abundance) [52], we provide a list of the most abundant proteins in UIP mucus (**Supplemental File 5**). We found that the mucus is enriched with immunoglobulins (Ig) which is in accord with increased protein expression of polymeric Ig receptor (PIGR) in the fibrotic honeycomb airway cells and mucus. In epithelial cells, PIGR mediates the transcytosis of Igs into the airway, which serves as a mucosal defence mechanism [53]. Given that fibrotic mucus is enriched with cellular infiltrates, we next focused our list using the ‘secretome’ (secreted proteins) dataset [54] and show that BPIFB1 was the most abundant secretome-associated protein found in fibrotic mucus whereas MUC5B is the ninth most abundant (**Figure 7A**; a full list in **Supplemental File 6**). This is consistent with BPIFB1 and MUC5B being amongst the most significantly expressed proteins in the fibrotic HC airway cells (**Figure 3A**). To validate some the most abundant protein hits, we show immunoreactivity for MUC5B, BPIFB1, PIGR, and TF within the UIP mucus plugs (**Figure 7B**). We also included the other gel-forming mucin, MUC5AC (60^th^ on the abundance list), which showed a patchy/incomplete staining pattern.

**Figure 7:**
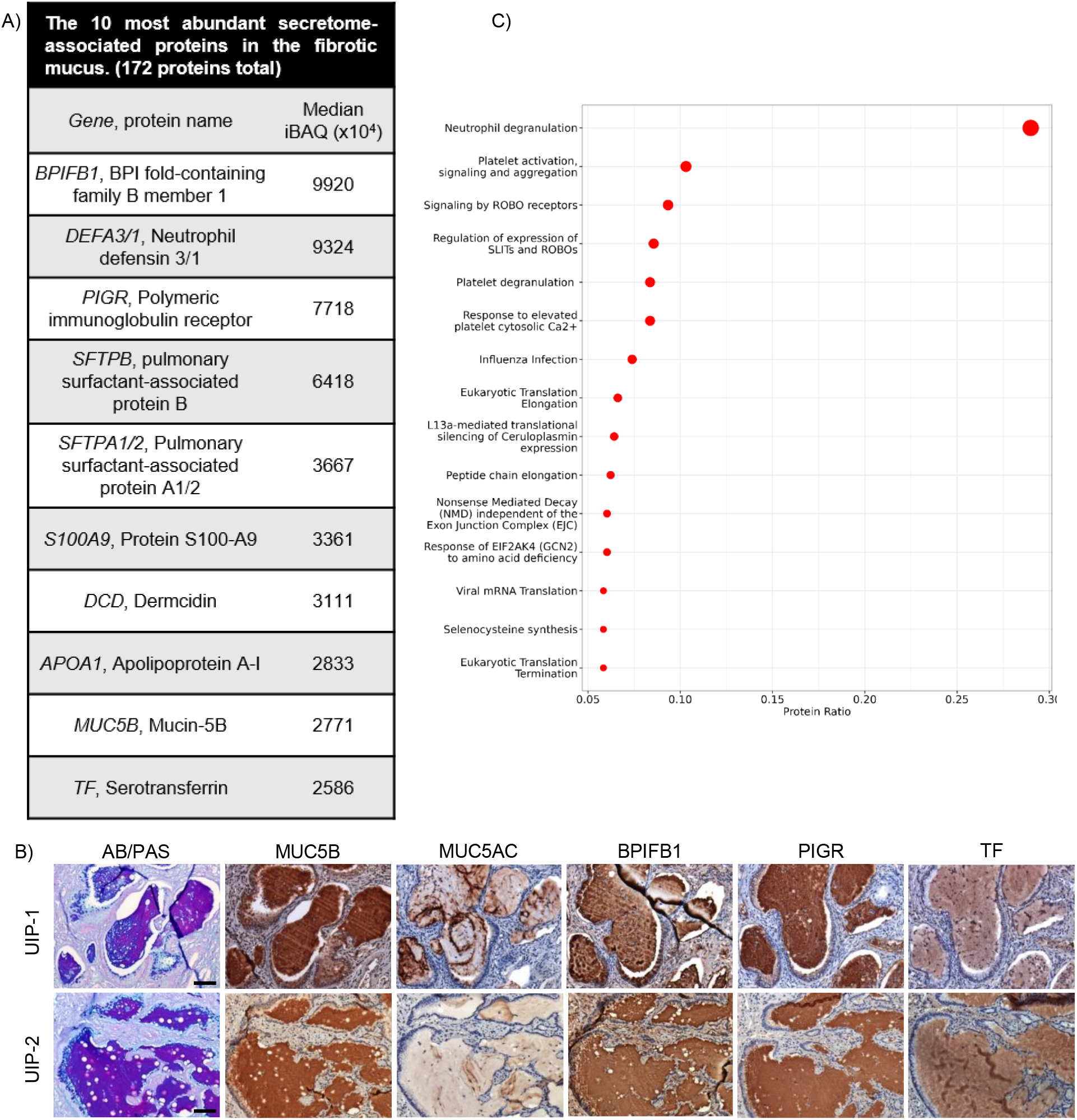
The composition of fibrotic mucus. Laser capture microdissection coupled mass spectrometry was performed on the mucus plugs of 6 Usual Interstitial Pneumonia (UIP) specimens. (**A**) A list of the most abundant secretome-associated proteins found in the fibrotic mucus shown as intensity Based Absolute Quantification (iBAQ). (**B**) Reactome pathway enrichment of UIP mucus represented as a dotplot. (**C**) Serial sections of UIP specimens stained for alcian blue/periodic acid Schiff’s (AB/PAS) or immunostained for MUC5B, MUC5AC, BPIFB1, PIGR, and TF (N = 4 UIP specimens with 2 representatives shown). Scale bar represents 100 microns.

Using the entire fibrotic mucus proteome, Reactome pathway enrichment analysis demonstrates that the mucus plug is defined by ‘neutrophil degranulation’ as the strongest category (**Figure 7C**). Neutrophil degranulation pathway is also implicated in SARS-CoV-2 lung infection models [55] and in chronic obstructive pulmonary disease (COPD) [56]; in IPF bronchoalveolar lavage fluid (BALF), proteins associated with neutrophil granules are amongst the most abundant [57].

### Fibrotic-derived lung mucus is distinct from cancer-derived lung mucus

To further understand mucus in the context of lung disease, we additionally performed LCM-MS on 6 mucinous adenocarcinoma (MA) specimens (**Supplemental Figure 2A**). MA is a lung cancer with pronounced mucus accumulation within the alveolar space [58, 59]. We detected a total of 535 proteins in MA mucus (found in 3 or more of the 6 samples; a full list in **Supplemental File 4**). Consistent with *MUC5AC* being the most abundantly expressed transcript in MA [60], MUC5AC protein is the second most abundant secretome-associated protein found in the mucus of MA (**Supplemental Figure 2B**, a full list in **Supplemental File 5**). Reactome pathway analysis demonstrates that MA mucus is defined by ‘Neutrophil degranulation’, like fibrotic mucus, as the strongest category (**Supplemental Figure 1C**).

**Supplemental Figure 2:**
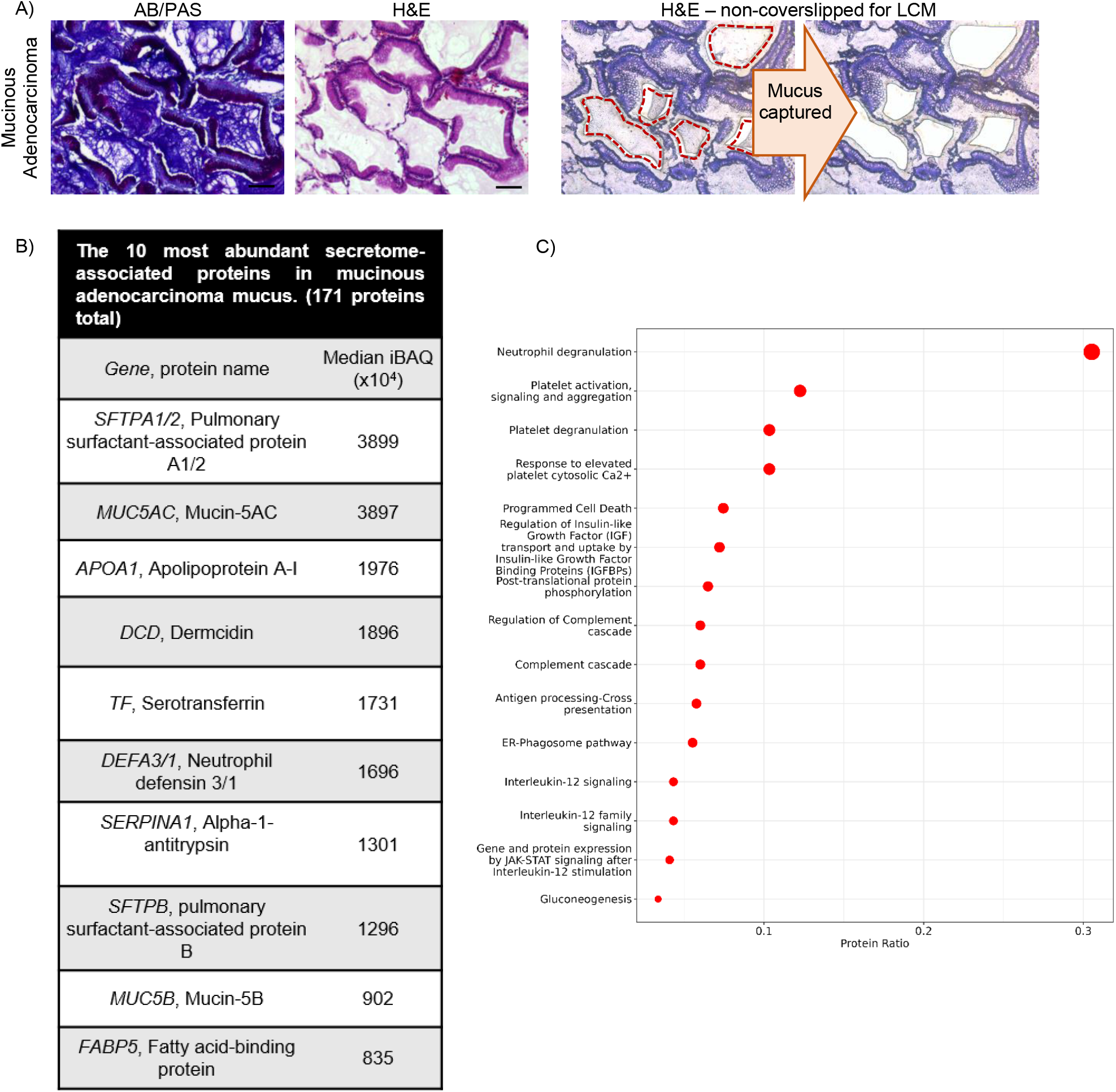
The mucus in mucinous adenocarcinoma (MA). (**A**) A MA specimen was serially sectioned at 5 microns and stained with alcian blue/periodic acid Schiff’s (AB/PAS) stain or Hemetoxylin & Eosin (H&E). Mucus was laser microdissected for mass spectrometry analysis. Scale bar represented 100 microns. (**B**) A list showing the most abundant secretome-associated proteins found in the mucus of MA. (**C**) Dotplot visualization of the MA mucus using Reactome pathway enrichment.

We next compared the mucus of MA to UIP. In total, we detected 707 lung mucus proteins (**Figure 8A**), with UIP having the most proteins detected (a full list in **Supplemental File 4**). A 3-D PCA analysis showed that UIP mucus samples largely cluster together, whereas only one MA sample overlaps with UIP (**Figure 8B**). Quantitative analysis of our data show that 9 proteins are significantly enriched in fibrotic mucus whereas 3 are significantly enriched in cancer (MA) mucus (**Figure 8C**, a full list in **Supplemental File 7**). To validate this result, we performed immunofluorescence on both MA (n = 3) and UIP specimens (n = 4) for MUC5B, MUC5AC, and BPIFB1 (**Figure 8D**). We found that UIP mucus has variable expression of MUC5AC (white arrows mark the absence of MUC5AC where MUC5B/BPIFB1 is present) in comparison to MUC5B. Note that one mucus plug was positive for MUC5AC, which resembles the chromogenic patchy/incomplete stain in **Figure 7B**; MUC5AC has been previously reported to have variable staining [10]. Inversely, MA mucus was positive for MUC5AC, whereas MUC5B and BPIFB1 are largely negative at the immunofluorescence level (**Figure 8E**, white arrows point where MUC5B/BPIFB1 are absent). Thus, a distinguishing factor for fibrotic/UIP mucus is the high abundance of MUC5B and BPIFB1, whereas MUC5AC is predominant in MA mucus.

**Figure 8:**
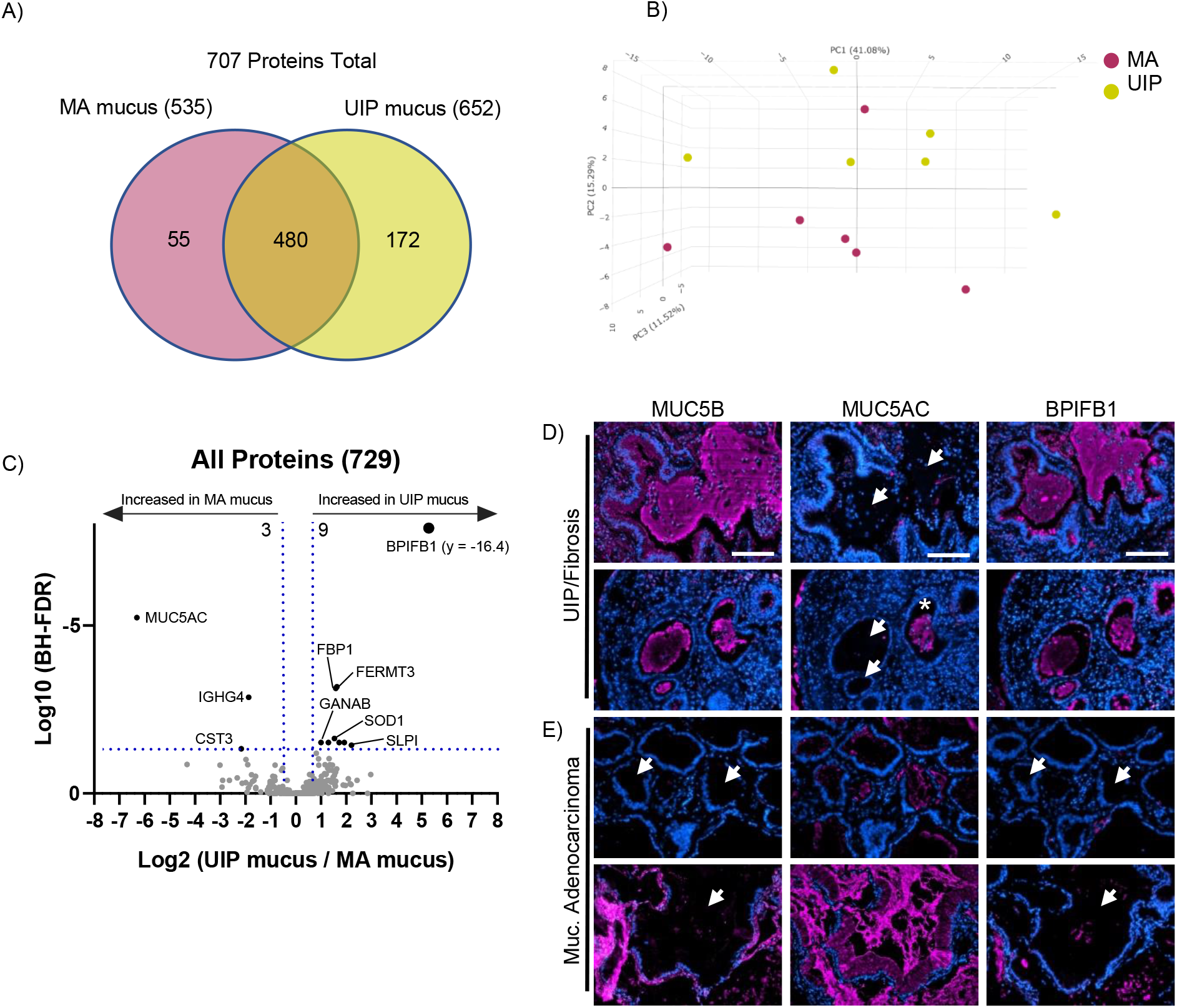
The mucus in usual interstitial pneumonia has distinct features from mucinous adenocarcinoma. (**A**) Venn diagram showing the proteins detected in the mucus of mucinous adenocarcinoma (MA) or usual interstitial pneumonia (UIP). (**B**) 3-dimensional principal component analysis for each mucus type. (**C**) Volcano plot comparing the UIP mucus to MA mucus showing the negative natural log of the false discovery values (FDR) values plotted against the base 2 log (fold change) for each protein. (**D**) Immunofluorescence for MUC5B, MUC5AC, and BPIFB1 in UIP mucus (n = 4 specimens) and MA mucus (n = 3 specimens) with representative images shown for each disease type. White arrows points to regions of mucus accumulation and asterisk shows positivity of MUC5AC within UIP mucus. Scale bar represents 100 microns.

## Discussion

In this work, we produced an unbiased spatial proteomic profile of the non-fibrotic, fibrotic uninvolved and honeycomb airway cells to create a tissue map that defines pathway changes along the progression of lung fibrosis (**Figure 9**). We showed that the structurally intact low-in-mucus airway cells in uninvolved regions of the fibrotic lung have an abnormal protein signature with increased Slits/Robo pathway as the strongest category; Slits/Robo pathway is also increased in the fibrotic HC airway cells. We confirmed that the fibrotic honeycomb airways are the site of mucin biogenesis with other categories related to protein modification and transport increased. Importantly, many proteins associated with ciliogenesis are decreased or absent from the fibrotic HC airways. In addition, honeycombing is associated with decreased extracellular matrix organization and elastic fibre formation. Lastly, the fibrotic mucus is enriched with immune defence proteins, including BPIFB1 and MUC5B, and is enriched with neutrophil degranulation pathway.

**Figure 9:**
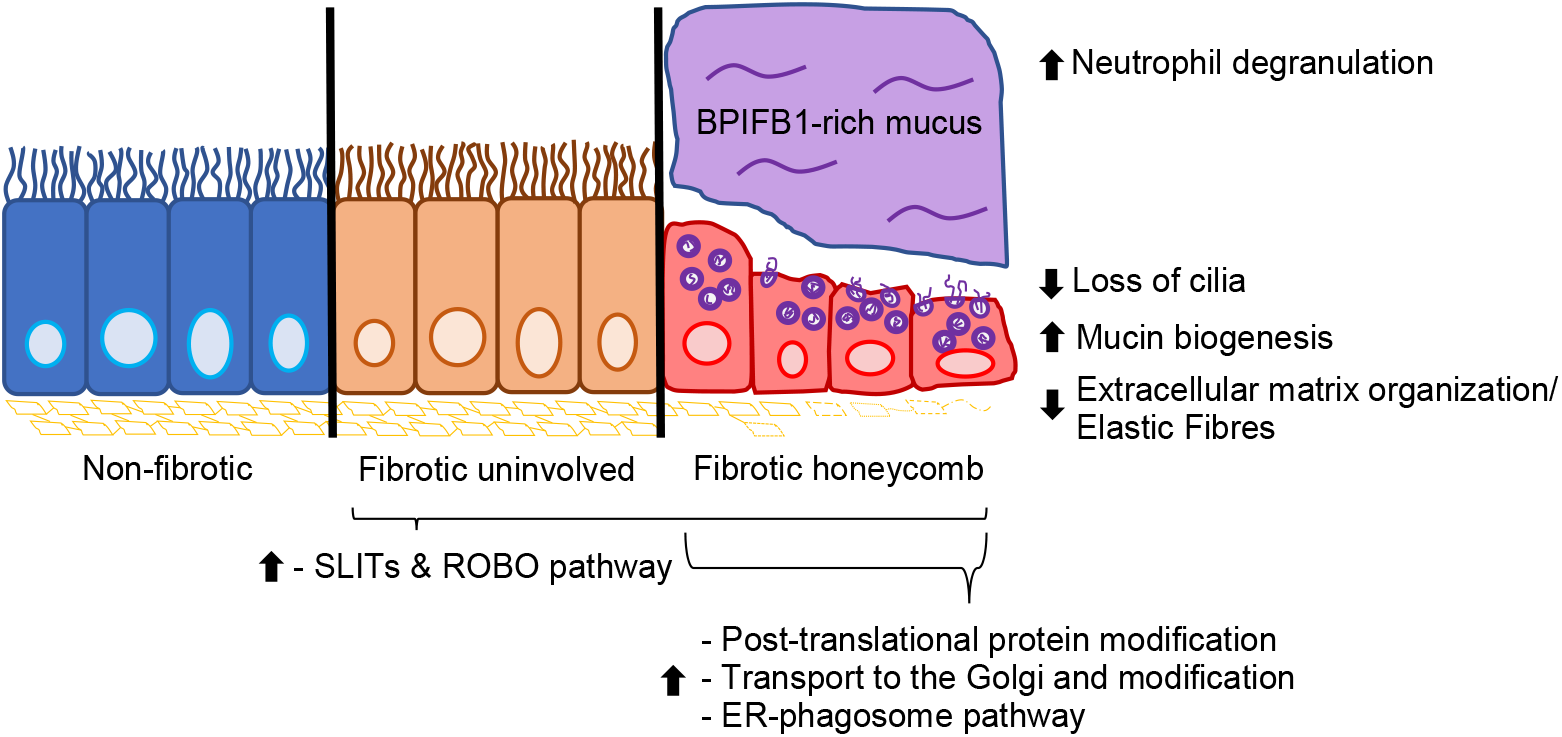
The fibrotic honeycomb airway. Spatial proteomics reveal that the fibrotic uninvolved airway cells (found in regions of structurally intact lung) have an abnormal protein signature. The fibrotic uninvolved airway cells, like the honeycomb (HC) airway cells (a pathological feature of lung fibrosis), are over-represented in proteins associated with SLITs and ROBO pathway. The fibrotic HC airway cells are further defined by increased pathways associated with mucin biogenesis, and a loss of both cilia and extracellular matrix organization/elastic fibres. We find that the mucus proteome is enriched with neutrophil degranulation pathway, with a marked increase of BPIFB1 protein.

Other groups support our results that demonstrate that the fibrotic uninvolved airway cells are abnormal at the structural and genetic level. Lung fibrosis is associated with a variety of genetic risk factors affecting epithelial cells, which may abnormally prime lung airway cells to fibrosis initiation [61]. Structurally, regions without microscopic fibrosis are shown to have reduced numbers of terminal airways and have an increase of airway wall areas [4–6], suggesting that early lung airway perturbations precede fibrotic extracellular matrix remodelling. Thus, it is plausible that airway cell dysfunction is an early event in the fibrotic process. Further LCM-MS studies with precise distance registration and patient genotyping will inform whether there exists a transition zone where a normal airway proteome is present, or perhaps it may be that the entire airway proteome is abnormal.

Aberrant ciliogenesis has previously been described in UIP/IPF. Whole transcriptomic studies demonstrate an elevation of cilium gene expression [62]. In contrast, our results demonstrate a reduced cilia-associated protein profile within the fibrotic HC airway cells. This likely reflects the advancement of our spatial proteomic capability or that previously observed increases in ciliary gene expression represent a compensatory mechanism occurring in response to loss of ciliated epithelial cells in fibrosis. Structurally, transmission electron microscopy demonstrates that the UIP/IPF distal airways display defects in microtubule organization, which will have detrimental effect on cilia function [39]. In the context in cystic fibrosis, it is reported that airway epithelial also have decreased ciliated cells with enhanced mucin expression [63]. Thus, future studies determining the mechanism of deranged ciliogenesis is warranted.

Current literature suggests that basal airways cells differentiate into either mucin producing cells or cilia-containing cells [64]. Our spatial proteomic data fits the notion that the HC airway microenvironment directs the differentiation of basal airway cells into mucin producing cells whereas the uninvolved airway microenvironment favors ciliated cells. Given that extracellular matrix governs cell differentiation and function [65], we speculate that changes to ECM properties (mechanical, composition, and topography) may play a role in airway cell differentiation and function. Prior work utilizing decellularized COPD airway tissue as a scaffold for cell-matrix interactions (as compared to donor) show that COPD matrix dramatically affects cilia gene expression in epithelial cells [66]. Other studies using decellularized UIP/IPF tissue confirm that fibrotic matrix is a driver of fibrosis progression [67]. Therefore, the changes in mucin and cilia-associated proteins may be reflective, or a consequence of the changes in airway ECM properties.

Our spatial proteomic data characterizing fibrotic HC airway cells (MUC5B-positive) are in agreement with sc-RNAseq data characterizing *MUC5B*-positive secretory cells in human lung. In one study, secretory airway cells have increased RNA expression of *MUC5B, LCN2, BPIFB1, SERPINB3, S100P, RARRES1, TSPAN8, CP*, and *FAM3D* [68], which are also increased or uniquely expressed at the protein level in fibrotic HC airway cells. Other mRNAs increased in *MUC5B*-positive secretory cells include *TSPAN1, AKR1C1, ZG16B, GSTA1*, and *SCGB1A1*, which are unchanged at the protein level in the fibrotic HC airway cells. A separate study showed that *MUC5B*-positive cells by sc-RNAseq have increased mRNA expression of *SCGB1A1, SCGB3A1, SLPI, BPIFB1, LCN2*, and *WFDC2* [69]. At the protein level, SLPI, BPIFB1, LCN2, and WFDC2 are increased or uniquely expressed in the fibrotic HC airway cells (SCGB1A1 and SCGB3A1 are unchanged at the protein level). Thus, the fibrotic HC airway cells represent a secretory cell phenotype. Future work integrating spatial multi-omic analysis (RNA and protein) will further our understanding of lung fibrosis.

To our knowledge, we are the first to determine the composition of UIP mucus plugs by using an LCM-MS approach. This approach allows precise capture of the entirety of mucus plugs without the introduction of contaminants (salivary and upper airway proteins) as seen by traditional BALF. Proteomic analysis of BALF (an unfixed or stained sample) from lung fibrosis patients show agreement with our findings. Several reports utilizing mass spectrometry approaches show increases of immunoglobulins, complement C3, transferrin, Apolipoprotein A1, plastin-2, annexin A2, and CCL18 in fibrotic lung BALF (summarized in [70]); all of which are detected in our LCM-MS dataset. In accord with our findings, Foster et al. demonstrated that MUC5B is an abundant protein in IPF BALF [57]. S100A9, detected by LCM-MS, is a potential BALF biomarker in IPF [71]. In addition, IPF patients with acute exacerbations show increased PIGR, LRG1, and SERPINA1 in BALF, which are also detected in our LCM-MS dataset [72]. Our LCM-MS approach is therefore a useful tool to determine the protein composition of mucus in archived FFPE specimens.

Our results demonstrate that the mucus found in lung cancer (mucinous adenocarcinoma; MA) has elevated levels of MUC5AC as compared to UIP mucus. A likely explanation is that the mucin-secreting cells comprising the UIP/IPF HC airway differ than the mucin-secreting cells in MA and/or that the environmental/immune signals controlling mucin production differ. For instance, reports show that there are 5X more MUC5B-positive cells versus MUC5AC-positive cells in the honeycomb airways of UIP/IPF, suggesting marked cell type heterogeneity [73]. In contrast, the morphology of MA cells are distinct and composed of goblet and/or columnar cells [74]. Another explanation is that *MUC5AC* gene expression is differentially regulated as compared to *MUC5B* [75]. For instance, *MUC5AC* gene expression is increased by IL-13. In other disease settings, *MUC5AC* mRNA is increased in asthma, whereas *MUC5B* levels are decreased [76]. Further studies determining the functional consequence of varying MUC5AC to MUC5B protein ratios on fibrosis progression are needed.

Increases of BPIFB1 in both the UIP mucus and HC airway cells is of interest. BPIFB1 is a secretory protein that is implicated in immune regulatory functions and shown to have anti-tumor effects (reviewed in [77]). In other lung disorders, BPIFB1 is increased in cystic fibrosis, COPD, asthma, and IPF [78]. It is decreased in nasopharyngeal carcinoma, gastic cancer, and lung cancer, which agrees with our findings that mucinous adenocarcinoma mucus has low expression of BPIFB1. Understanding of its function in lung fibrosis is currently incomplete.

## Conclusion

Spatial proteomics has allowed us to create an unbiased protein tissue map of the fibrotic/UIP lung airway cells. We show that the fibrotic honeycomb airway cells are the active site of mucin biogenesis affiliated with a loss of cilia. Importantly, we show that the fibrotic uninvolved airway cells have an abnormal protein signature. Therapeutic intervention of the fibrotic uninvolved airway cells may therefore slow fibrosis progression.

## Materials and Methods

### Histological staining

Five micron sections of formalin-fixed and paraffin-embedded (FFPE) specimens were H&E-stained by using an automated stainer (Leica XL) at the University of Manchester Histology Core Facility as previously described [19]. Importantly, slides were stored at 4oC for up to one week while laser capture microdissection (LCM) was being performed. Captured material was stored at −80°C until all samples were ready for mass spectrometry processing. Alcian Blue/Periodic Acid Schiff (AB-PAS) was performed as follows. De-paraffinized slide sections were incubated for 5 minutes in 1% alcian blue 8GX (Sigma; A5268), 3% acetic acid. Slides were then washed in tap water followed by a 5-minute incubation in 1% periodic acid (Sigma; 375810). Finally, slides were washed in tap water and incubated in Schiff’s reagent (Sigma – 3952016) for 15 minutes. After extensive washing in tap water, slides were coverslipped without counterstain. For pentachrome, we followed a protocol as previously described [19].

For immunohistochemistry (IHC), we utilized the Novolink Polymer Detection Systems (Leica, RE7200-CE) as previously described in detail [79]. We used the following antibodies anti-BPIFB1 (Abcam; ab219098, titre 1:60,000), anti-elastin (Proteintech; 15257-1-AP; titre 1:16,000) anti-PIGR (Abcam; ab224086, titre1:8,000), anti-serotransferrin (Abcam; ab268117, titre 1: 30,000). Anti-MUC5B (titre 1:10,000) and anti-MUC5AC (titre 1:12,000) was previously purified and used here [80]. For all samples, we used antigen heat retrieval using citrate buffer pH 6.0 (Sigma, C9999), with the exception of EDTA pH 9.0 antigen heat retrieval for serotransferrin and elastin. Slides were hematoxylin counterstained and coverslipped using permount (ThermoScientific, SP15). For MUC5B, immunostains followed a modified protocol. After citrate buffer pH 6.0 antigen heat retrieval, the sections underwent reduction and alkylation. Sections were reduced by incubation at 37oC for 30 minutes in 10 mM DTT, 0.1 M Tris/HCl pH 8.0. Sections are washed in water and then incubated in 25 mM Iodoacetamide, and 0.1 M Tris/HCl pH 8.0 for 30 minutes at room temperature (kept in the dark). Lastly, sections were washed in water followed by blocking and primary antibody incubation.

For immunofluorescence, dewaxed slides were subjected to citrate buffer pH 6.0 antigen heat retrieval and probed overnight with anti-MUC5AC (titre 1:100), anti-MUC5B (post reduction/alkylation; titre 1:100), or BPIFB1 (Abcam; ab219098, titre 1:100). Sections were then incubated with secondary anti-mouse fluorophore 680 (Invitrogen, A21058, 1:500) or anti-rabbit fluorophore 680 (Invitrogen; A21109; 1:500) for 1 hour. Sections were coverslipped using ProLong antifade with DAPI (Invitrogen; P36931).

### Laser Capture Microdissection

The MMI CellCut Laser Microdissection System (Molecular Machines & Industries) was used to capture regions of interest on MMI membrane slides (MMI, 50102) as previously described [19, 20]. For this set of experiments, we collected a volume 0.03 mm^3^ of tissue per sample.

### Histological Imaging

For fluorescence microscopy, all stains were performed at the same time. In addition, images were taken at the same intensity utilizing EVOS FL imaging system (ThermoScientific). For light microscopy, we used a DMC2900 Leica instrument with Leica Application Suite X software.

## Data Availability

We have deposited the raw mass spectrometry data files to ProteomeXchange under the identifier of PD036465.

### Mass spectrometry sample preparation

Samples were prepared as described [19, 20]. In short, samples underwent a series of steps to maximize protein yield, including high detergent treatment, heating, and physical disruption.

### Liquid chromatography coupled tandem mass spectrometry

The separation was performed on a Thermo RSLC system (ThermoFisher) consisting of a NCP3200RS nano pump, WPS3000TPS autosampler and TCC3000RS column oven configured with buffer A as 0.1% formic acid in water and buffer B as 0.1% formic acid in acetonitrile. An injection volume of 4 µl was loaded into the end of a 5 µl loop and reversed flushed on to the analytical column (Waters nanoEase M/Z Peptide CSH C18 Column, 130Å, 1.7 µm, 75 µm X 250 mm) kept at 35 °C at a flow rate of 300 ηl/min with an initial pulse of 500 ηl/min for 0.1 minute to rapidly re-pressurize the column. The separation consisted of a multistage gradient of 1% B to 6% B over 2 minutes, 6% B to 18% B over 44 minutes, 18% B to 29% B over 7 minutes and 29% B to 65% B over 1 minute before washing for 4 minutes at 65% B and dropping down to 2% B in 1 minute. The complete method time was 85 minutes.

The analytical column was connected to a Thermo Exploris 480 mass spectrometry system via a Thermo nanospray Flex Ion source via a 20 µm ID fused silica capillary. The capillary was connected to a fused silica spray tip with an outer diameter of 360 µm, an inner diameter of 20 µm, a tip orifice of 10 µm and a length of 63.5 mm (New Objective Silica Tip FS360-20-10-N-20-6.35CT) via a butt-to-butt connection in a steel union using a custom-made gold frit (Agar Scientific AGG2440A) to provide the electrical connection. The nanospray voltage was set at 1900 V and the ion transfer tube temperature set to 275 °C.

Data was acquired in a data dependent manner using a fixed cycle time of 1.5 sec, an expected peak width of 15 sec and a default charge state of 2. Full MS data was acquired in positive mode over a scan range of 300 to 1750 Th, with a resolution of 120,000, a normalized AGC target of 300% and a max fill time of 25 mS for a single microscan. Fragmentation data was obtained from signals with a charge state of +2 or +3 and an intensity over 5,000 and they were dynamically excluded from further analysis for a period of 15 sec after a single acquisition within a 10-ppm window. Fragmentation spectra were acquired with a resolution of 15,000 with a normalized collision energy of 30%, a normalized AGC target of 300%, first mass of 110 Th and a max fill time of 25 mS for a single microscan. All data was collected in profile mode.

### Mass spectrometry data analysis and statistics

Raw data for regional samples were processed using MaxQuant [81] version 1.6.17.0 against the human proteome obtained from uniprot (May 2021) [82]. Raw data for UIP and MA mucus samples were processed using MaxQuant [81] version 2.0.3.0 against the human proteome obtained from uniprot (May 2022) [82]. All Maxquant processing were performed with a fixed modification of carbamidomethylation of cysteine, with variable modifications of methionine oxidation and protein N-terminal acetylation. Precursor tolerance was set at 20ppm and 4.5pm for the first and main searches, with MS/MS tolerance set at 20ppm. A false discovery rate (FDR) of 0.01 was set for PSM and protein level, up to two missed cleavages were permitted and “match-between-runs” was selected.

Stastical analysis was carried out in R (v4.1.2) [83] using the MSqRob package (v0.7.7) [84]. Significantly changing proteins were taken at a 5% false discovery rate (FDR). Pathway analysis utilising Reactome Pathways was performed on significantly changing proteins using the R package ReactomePA (1.38.0) [85]

### Study Approval

For this study, we utilized a variety of Research Ethics Committee (REC) protocols to obtain patient-consented lung tissue: REC#14/NW/0260 (provided by JFB and RVV, Manchester, United Kingdom) for transplanted fibrotic lung; REC#20/NW/0302 (provided by MAM and FG, Manchester, United Kingdom) for non-fibrotic lung specimens; REC#18/NW/0092 (provided by Manchester Cancer Research Centre Biobank, Manchester, United Kingdom) for mucinous lung adenocarcinoma. Usual Interstitial Pneumonia (UIP) specimens were defined by current guidelines [1, 86]. Non-fibrotic controls were collected from morphologically normal lung tissue distal to tumor during resection (fibrotic and control patient demographics may be found in **Supplemental Figure 3**). Mucinous adenocarcinoma (MA) was defined by current guidelines (MA patient demographics may be found in **Supplemental Figure 4**) [59]. In this study, we utilized 10 UIP, 6 non-fibrotic, and 6 MA specimens.

## Author Contributions

J.A.H. designed and conducted all LCM-MS experiments and C.L. performed the associated analyses. L.D. and S.P. performed the immunohistochemistry/immunofluorescence and imaging. M.A.M. assisted in characterizing the histological stains and identification of clinical morphologies associated with Usual Interstitial Pneumonia and Mucinous Adenocarcinoma. R.V.V., J.F.B., M.A.M. and F.G. contributed to reagents. J.A.H. wrote the manuscript with all author inputs. J.A.H. and D.J.T conceived and supervised the project.

## Acknowledgements

The authors would like to thank the Histology, BioMS, and BioImaging Facilities at University of Manchester for making this work possible. The authors would like to acknowledge the Manchester Allergy, Respiratory and Thoracic Surgery (ManARTS) Biobank; Manchester Cancer Research Centre (MCRC) Biobank; Transplant Unit staff at University Hospital of South Manchester NHS Foundation Trust; and the North West Lung Centre Charity (NWLC) for supporting this project. The views expressed in this publication are those of the authors and not necessarily those of ManARTs, MCRC, NHS, or NWLC. In addition, we thank the study participants for their contribution.

